# SMART platform for enzyme engineering: A novel single-molecule display approach for evolution and functional studies of D-amino acid oxidase

**DOI:** 10.1101/2025.03.23.644798

**Authors:** Kalhari Munaweera, Nana Odake, Kakeru Ikeda, Bo Zhu, Maurizio Camagna, Tomokazu Ito, Tetsuya Kitaguchi, Naoto Nemoto, Hideo Nakano, Jasmina Damnjanović

## Abstract

This study introduces SMART (<u>S</u>ingle <u>M</u>olecule <u>A</u>ssay on <u>R</u>ibonucleic Acid by <u>T</u>ranslated Product), an innovative *in vitro* platform integrating mRNA display, next-generation sequencing, and bioinformatics to address several key challenges in enzyme engineering. The characteristic feature of SMART is its auxiliary unit, attached to the mRNA-displayed enzyme library to mediate the biotinylation of reactive enzyme variants and enable their enrichment by streptavidin beads pull-down. The unit comprises a hairpin single-stranded DNA that hybridizes with the mRNA and anchors a DNA-binding protein, single-chain Cro, which carries functional auxiliary molecules specific to the enzyme of interest. Here, we report on the establishment of SMART for the engineering of oxidases, specifically D-amino acid oxidase (DAAO) from *Schizosaccharomyces pombe*, with engineered ascorbate peroxidase 2 as its auxiliary enzyme. In a single selection round of a DAAO library with randomized catalytic residue Y232 and D-Alanine as the substrate, we identified several catalytically active enzyme variants with altered substrate specificity in a matter of days. This surpasses traditional enzyme engineering methods in speed, cost, library capacity, and precision. This research underscores the potential of SMART for the engineering of various oxidases, and other enzymes after the corresponding adjustment of the auxiliary unit.

## INTRODUCTION

Biocatalysis gained enormous importance in all aspects of industry^[1]^. To become useful biocatalysts, enzymes are developed by protein engineering (i.e. enzyme engineering). Directed evolution is a proven strategy for enzyme engineering that mimics natural selection in a test tube^[2]^. Selection is performed *in vivo* or *in vitro* with iterative rounds of mutagenesis and selection to obtain improved enzyme variants^[3]^. While mutagenesis can be simply done by polymerase chain reaction (PCR) with mutagenic primers using universal and well-established methods, selection must be enzyme-specific^[4]^ and is considered the bottleneck in terms of time, cost, efficiency, and labour.

In protein engineering by directed evolution or semi-rational design, *in vivo* selection methods face challenges stemming from the use of cells, mainly library size (up to 10^9^ variants/mL) limited by the cell transformation efficiency, and bias toward well-expressed, non-toxic proteins. In contrast, *in vitro* selection methods, such as ribosome display^[5]^, mRNA display^[6]^, liposome display^[7]^, and *in vitro* compartmentalization (IVC) based on microdroplet technology^[8]^ use cell-free protein synthesis (CFPS) with genotype-phenotype link realized via oligonucleotide linkers or compartmentalization. These methods have tackled the limitations of *in vivo* methods by eliminating host-related biases^[9]^, and by enabling time and cost-effective operation, as well as the use of larger library size (about 10^12-14^ variants/ml).

Despite the advantages of *in vitro* methods, their routine application in enzyme engineering has yet to reach its full potential. Ribosome display has been used for a few proof-of-concept selections of dihydrofolate reductase, β-lactamase, DNA ligase, and sortase^[10]^. In a pioneering work by the Seelig group, a completely artificial ligase has been selected using mRNA display^[11]^, strongly demonstrating the power of *in vitro* methods in enzyme engineering. Proof-of-concept selections of methyl transferase, phosphotriesterase, and a restriction nuclease have been done by IVC coupled with fluorescence-activated droplet sorting^[12]^. A notable example of the IVC approach includes microbead display, where sensitive ultra-high-throughput screening was established for horseradish peroxidase (HRP)^[13]^. Overall, past experiences point out the difficulties in establishing functional genotype-phenotype linkage, the design of reliable selection conditions, robustness of protocols, and cost performance (mainly equipment cost) as the key factors that have limited the use of *in vitro* methods in enzyme engineering.

To address these challenges, we developed a novel *in vitro* platform <u>S</u>ingle <u>M</u>olecule <u>A</u>ssay on <u>R</u>ibonucleic acid by <u>T</u>ranslated product (SMART) which integrates mRNA display, next-generation sequencing (NGS), and bioinformatics (Scheme 1). The SMART platform enables rapid (4-5 hours by a single person), and cost-effective selection of enzymes from large combinatorial libraries with precise control over selection conditions. The library diversity is introduced at the DNA level, with mutagenesis of key enzyme amino acid positions identified through structural and ancestral trace analysis. Following *in vitro* transcription, the mRNA library is chemically linked to a puromycin linker^[14]^ (Fig. S1), for covalent binding of each *in vitro* translated enzyme to its corresponding mRNA.

The auxiliary unit, a characteristic feature of the SMART platform, is added to the mRNA-displayed enzyme under mild conditions. This unit consists of a hairpin DNA containing a segment that hybridizes with a complementary mRNA region (~20 bases) providing the controlled immobilization of the auxiliary unit, and a hairpin region where the DNA-binding protein single-chain Cro (scCro) binds with high affinity (K_D_ ~ 14 pM)^[15]^. Various functional molecules, such as auxiliary enzymes or enzyme substrates, can be attached to scCro^[13]^, creating an auxiliary unit tailored for a specific enzyme selection. Following the selection step, enzymes with desired characteristics are separated using streptavidin beads pull-down of biotinylated single-molecule display complexes. Recovered mRNA is reverse transcribed and resulting cDNA is amplified by PCR for preparation of the DNA library used in the following selection round and for the NGS analysis. This study describes the establishment of SMART platform for engineering of oxidases, namely a flavoenzyme D-amino acid oxidase (DAAO) and its evaluation through selection of active DAAO variants from a small mutagenic library targeting one of the active site amino acid positions. We have chosen DAAO for its high potential in quantitative measurement of individual D-amino acids recognized as health biomarkers in human blood and urine^[16]^.

## RESULTS AND DISCUSSION

To make DAAO fit for this application, wild-type (WT) DAAO^[17]^ needs to be engineered to modify its substrate specificity and improve its stability. The DAAO gene from the yeast *Schizosaccharomyces pombe* (SpDAAO) encodes a protein of 348 amino acids which forms nonessential homodimers, like most of the other homologous DAAO genes ^[17, 18]^. We hypothesized that the use of the dimeric enzyme form in SMART could be a source of false positives through heterodimer formation (Fig. S2). To ensure consistent and interpretable outcomes, we first prepared a monomeric functional form of SpDAAO by rational design. First, the model structure of dimeric WT SpDAAO was prepared by Swiss-Model server (https://swissmodel.expasy.org), and based on interface interactions, E49I, W253I, and R255L (Fig. S3A), mutations were chosen and individually introduced into the SpDAAO gene, aiming to disrupt interactions at the dimer interface. R255L SpDAAO was shown to form a monomeric structure (Fig. 1A left panel, S3B, S3C) and exhibited the highest enzymatic activity in comparison to the other two variants (Fig. S4A) while retaining a similar expression level (Fig. S4B). In comparison to the WT SpDAAO, the R255L variant showed overall moderately lower activity (Fig.1C).

**Figure 1.**
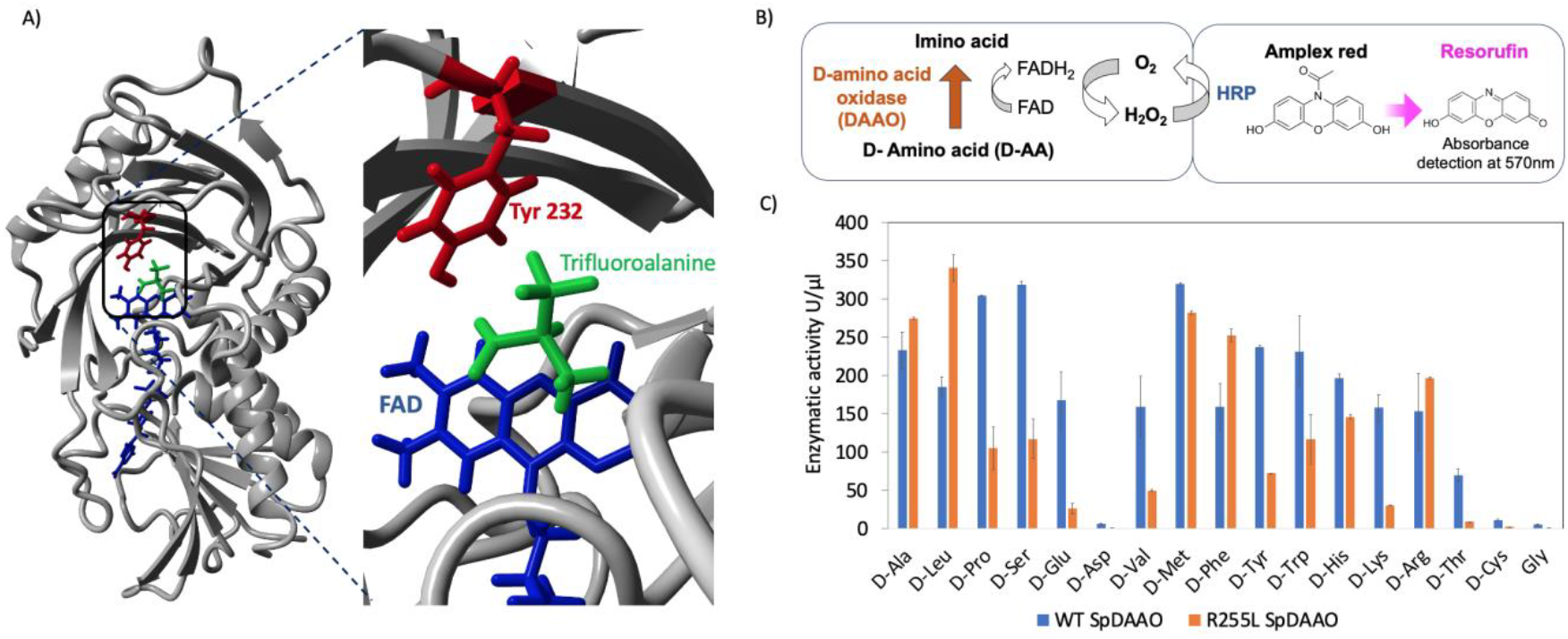
A) The model structure of R255L SpDAAO obtained by Yasara *Structure* using dimeric model as a template, with bound flavin adenine dinucleotide (FAD; blue), trifluoroalanine (green) and Y232 (red). B) The concept of Amplex red assay showing the coupled DAAO-HRP reaction. H_2_O_2_ produced in the DAAO reaction is used by HRP to convert Amplex red into resorufin. C) The reactivity of WT and R255L SpDAAO assessed using the Amplex red assay against a variety of D-amino acids (error bars indicate S.D. of two independent measurements).

As a functional molecule of the auxiliary unit for SpDAAO selection, we chose engineered ascorbate peroxidase 2 (APEX2)^[49]^ (predicted single-molecule display structure shown in Fig. S5). APEX2 biotinylates the proximal tyrosine-containing biomolecules rapidly at a distance of 1-10 nm, ensuring that only biomolecules closest to the source of H_2_O_2_ are labelled^[19]^. To anchor APEX2, after testing different options, we successfully implemented an ORC hairpin DNA (Fig. S6) which binds scCro and holds the APEX2-scCro fusion protein (Fig. S7) sufficiently close to displayed SpDAAO, ensuring efficient tandem reaction (Fig. S8) between the reactive SpDAAO variants and APEX2.

Initially, we validated SMART by checking the photocrosslinking of the mRNA, and the formation of functional mRNA display with the monomeric R255L SpDAAO (Figs. S9, S10A). We next performed a proof-of-concept SMART selection of 1:1, 1:10, and 1:100 model libraries composed of indicated molar ratios of active and inactive SpDAAO genes respectively, with D-Alanine (D-Ala) as the substrate. The inactive SpDAAO used in this study was R255L/Y232A double mutant. Since Tyr at position 232 of SpDAAO has a known role in substrate binding^[20]^ and is highly conserved among the predicted ancestral proteins (Fig. S11) and existing DAAO homologs ^[17a, 18a]^ (Fig. S12), we hypothesized that Y232A mutation will render the enzyme inactive. Indeed, Y232A/R255L lost its activity against D-Ala (Fig. S10B). Additionally, we introduced the *Xho*I restriction site into the gene of the double mutant, which is absent in the R255L SpDAAO (Fig. S13A), to analyze the composition of the enriched and remaining pools after the restriction digestion. After a single SMART selection round, active R255L SpDAAO was successfully enriched, while inactive genes corresponding to the Y232A/R255L variant were retained in the unselected pool (Fig. S13B-V). This result represents the first proof-of-concept of the new SMART platform and prompted further experiments to prove its efficacy.

A small library was prepared by site-saturation mutagenesis at position 232 of R255L (Y232X/R255L; Fig. 2A-I) and WT (Y232X WT; Fig. 2A-II) SpDAAO gene in anticipation of Y232 enrichment if the selection proceeds as designed. We included WT SpDAAO in this experiment to investigate the possible difference in selection efficiency between monomeric and dimeric enzyme libraries. After confirming DNA, mRNA, mRNA-linker, and mRNA display library preparation (Fig. 2B), the activity-based selection was performed using D-Ala as the substrate, with a reaction time of 10 minutes at 37°C. The quality of selection was first assessed by a bulk activity assay of input and enriched library DNA pools, after the corresponding DNA recovery and CFPS using PURE*frex* 2.1 (Gene Frontier; Fig. 2C).

**Figure 2.**
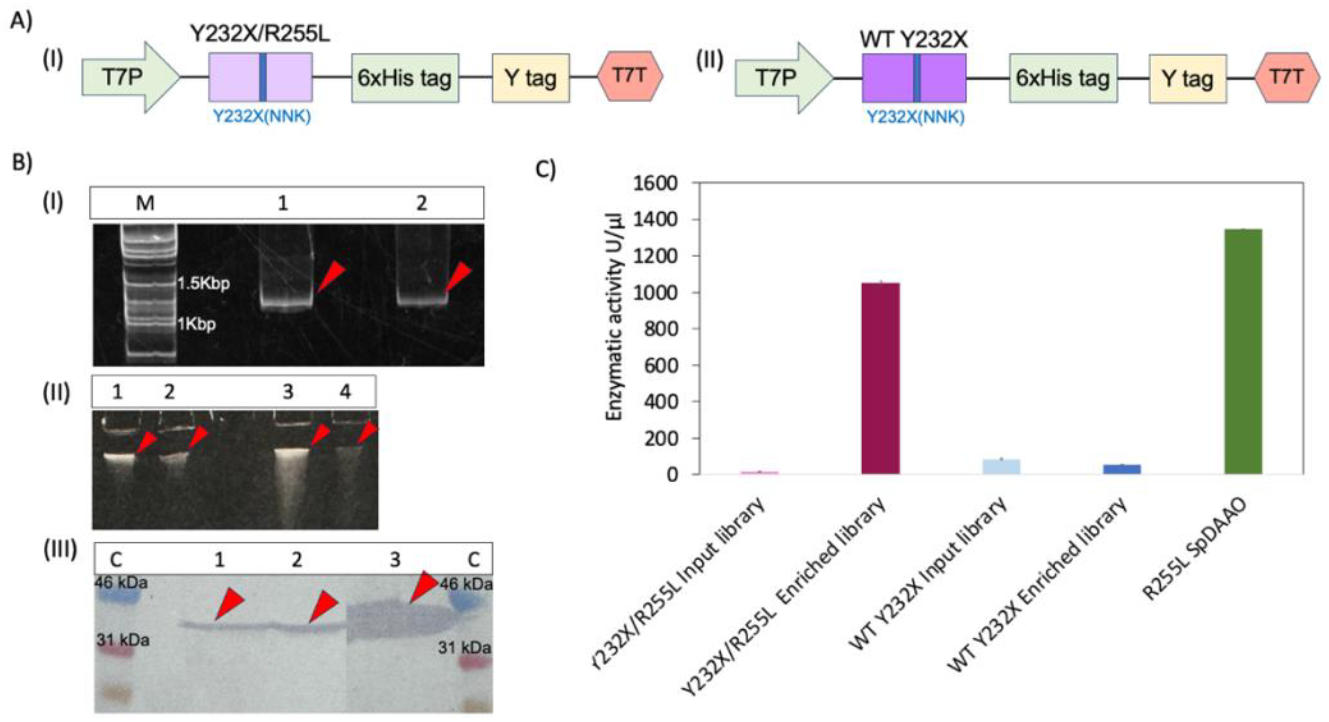
Evaluation of WT Y232X and Y232X/R255L libraries of SpDAAO. A) (I) The outline of Y232X/R255L SpDAAO DNA library; (II) The outline of WT Y232X SpDAAO DNA library; each containing a T7 promoter, T7 terminator, 6xHis tag, and a Y tag incorporating the sequence for mRNA hybridization to the puromycin linker. B) (I) DNA-PAGE of Y232X/R255L and WT Y232X DNA libraries. Lane M: Excel Band 1kb DNA Ladder, Lane 1: WT Y232X DNA library, Lane 2: Y232X/R255L DNA library; (II) Urea-PAGE of mRNA libraries before and after photocrosslinking stained with SYBRGold. Lane 1: WT Y232X mRNA library after photocrosslinking, Lane 2: WT Y232X mRNA library before photocrosslinking, Lane 3: Y232X/R255L mRNA library after photocrosslinking, Lane 4: Y232X/R255L mRNA library before photocrosslinking; (III) Western blot detected using Anti-His-tag Monoclonal Antibody HRP-DirecT, stained with TMB solution. Lane C: Protein MultiColor, Stable II, DynaMarker, Lane 1: WT Y232X mRNA display library after mRNA digestion with RNaseA, Lane 2: Y232X/R255L mRNA display library after mRNA digestion with RNaseA, Lane 3: APEX2-scCro synthesized by CFPS. C) 4-Aminoantipyrene assay of SpDAAO variants in input and enriched libraries after CFPS.

The results showed 100-fold higher activity of the Y232X/R255L SpDAAO enriched library in comparison with that of the input library. This trend was not observed for the WT Y232X SpDAAO enriched library which showed similar low activity as its input library. This result indicated that the quality of selection is heavily affected by the dimeric nature of the displayed enzyme. To analyze the composition of mutations in the input and enriched libraries, we carried out NGS using the NextSeq550 Illumina platform, with DNA prepared by PCR amplification of input and enriched library cDNA pools using designated primer pairs (Table S2). The Y232X/R255L library showed a high enrichment of Y, V, F, S, R, D, W, I, and P sequences (Table 1). As anticipated, the WT Y sequence was among the enriched variants, indicating that the SMART platform functions as per its design. In contrast, we found that there was no true enrichment of any sequence from the WT Y232X SpDAAO library (Table S1), indicating that the dimeric conformation is not suitable for selection using the current version of the SMART platform.

**Table 1.** Count numbers and enrichment factors of SpDAAO variants in Y232X/R255L input and enriched libraries. Positive enrichment score: EF>1, Negative enrichment score: EF <1.

| Residue at 232 position | Input library counts | Enriched library counts | Enrichment factor |
| --- | --- | --- | --- |
| Y | 51234 | 1011468 | 20.4 |
| V | 2296 | 112658 | 50.7 |
| F | 28106 | 104563 | 3.8 |
| S | 3054 | 97912 | 33.1 |
| R | 405 | 75918 | 193.6 |
| D | 1533 | 32114 | 21.6 |
| W | 1339 | 25240 | 19.5 |
| I | 192 | 18905 | 101.7 |
| P | 235 | 16855 | 74.1 |
| N | 122 | 479 | 4.1 |
| H | 17 | 302 | 18.4 |
| A | 1909039 | 998402 | 0.5 |
| G | 175621 | 64912 | 0.4 |
| C | 61378 | 47658 | 0.8 |
| L | 328410 | 1982 | 0.0 |
| K | 1304 | 1030 | 0.8 |
| T | 132821 | 769 | 0.0 |
| E | 177 | 98 | 0.6 |
| M | 209 | 31 | 0.2 |
| Total counts | 2697497 | 2611301 |  |

We next proceeded to assay the Y232X/R255L SpDAAO variants for their enzymatic activity. Genes of several variants with positive and negative enrichment scores were prepared and used as templates for the synthesis of the corresponding proteins by CFPS (Fig. 3A). The activities of CFPS-made enzymes were evaluated by Amplex red assay against D-Ala (Fig. 3B).

**Scheme 1.**
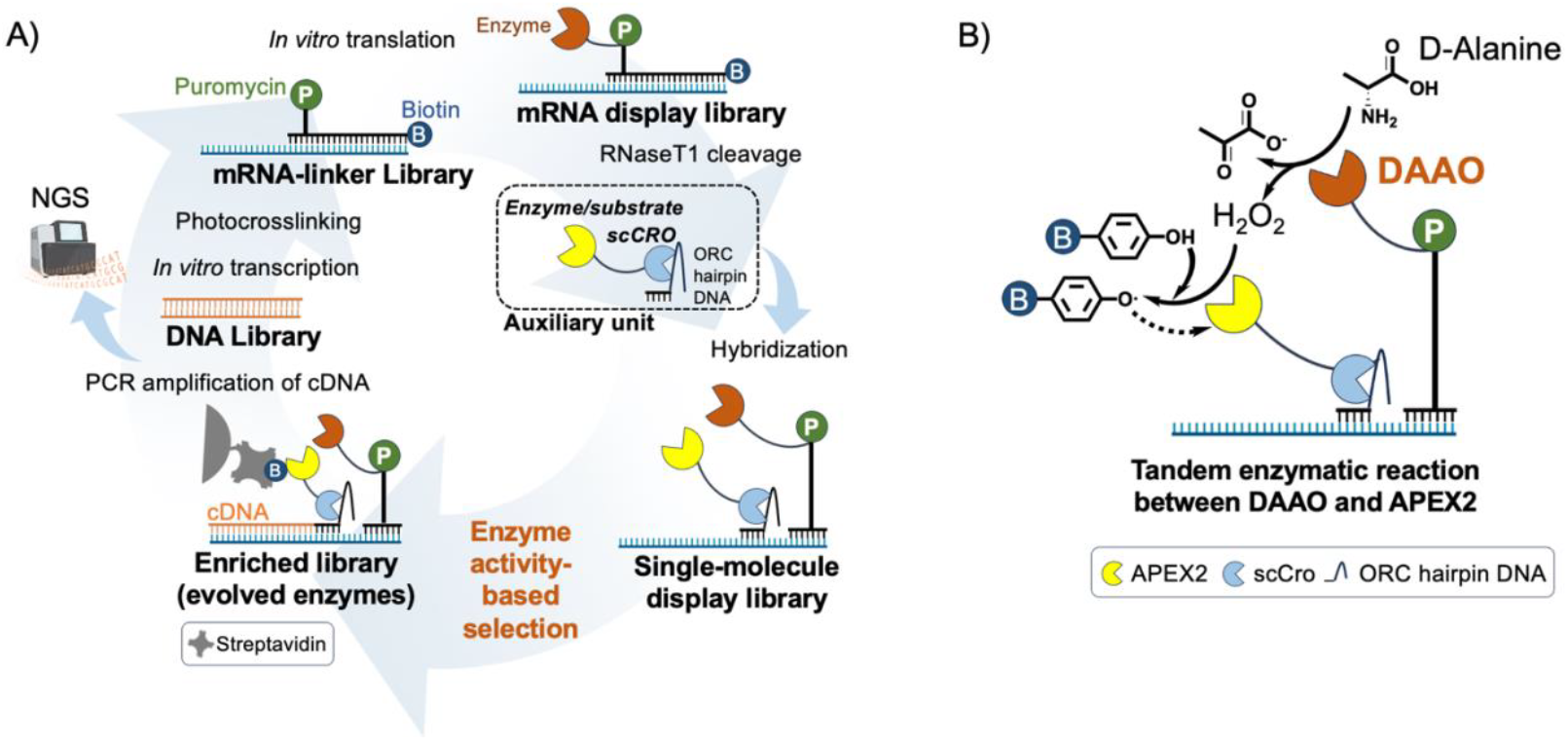
A) Outline of the SMART platform. After DNA library preparation, *in vitro* transcription, photocrosslinking, and *in vitro* translation, the auxiliary unit is attached to the display complex. The resulting single-molecule display library is subjected to enzyme activity-based selection resulting in biotinylation of complexes with active displayed enzymes, which can be captured by streptavidin beads pull-down. Input (original) and enriched library are then used in reverse transcription and PCR in preparation for NGS. PCR-amplified enriched library is also used to prepare DNA for the next selection round. B) Principle of DAAO selection with D-Ala as substrate. H_2_O_2_ generated by D-Ala oxidation is detected by APEX2, and subsequently used for conversion of biotin-tyramide into biotinyl free radical for proximity labelling of the complexes containing active DAAO variants. DAAO activity-dependent biotinylation serves to separate the evolved enzyme variants from the rest of the library using streptavidin beads pull-down.

**Figure 3.**
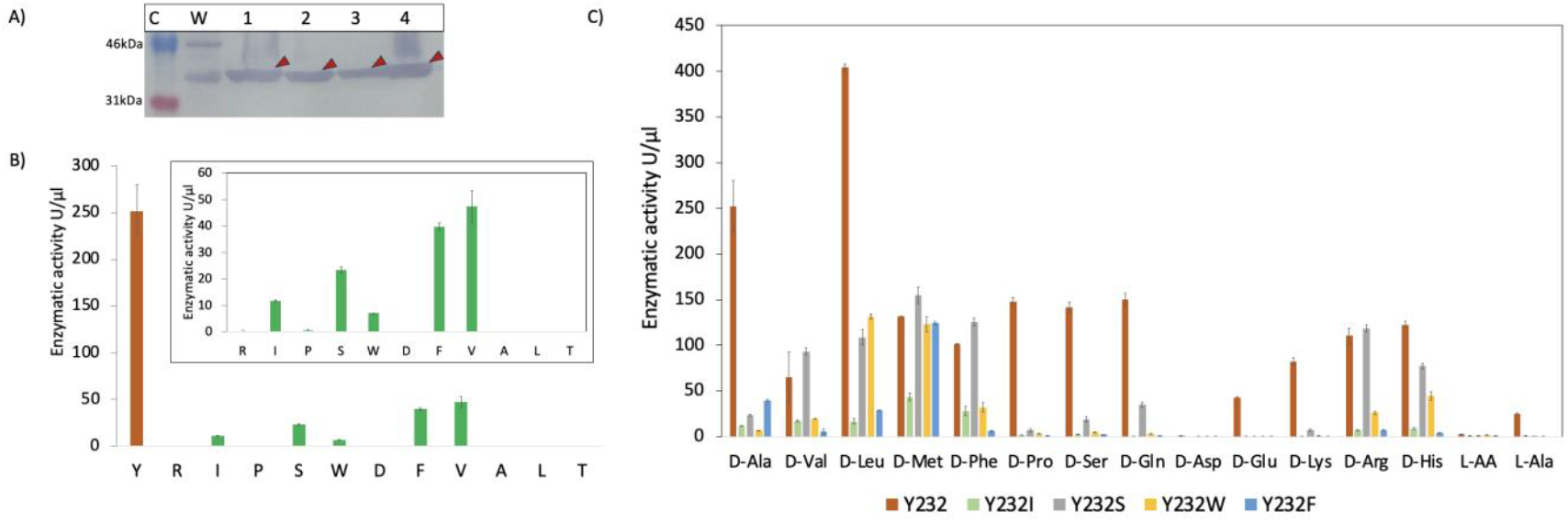
Characterization of CFPS-made SpDAAO variants enriched from Y232X/R255L library. A) Western blot of SpDAAO variants detected using Anti-His-tag Monoclonal Antibody HRP-DirecT, stained with TMB solution. Lane C: Protein MultiColor, Stable II, DynaMarker, Lane W: Dr. Western marker, Lane 1: WT SpDAAO, Lane 2: R255L SpDAAO, Lane 3: Y232A/R255L SpDAAO, Lane 4: Y232R/R255L SpDAAO. B) Amplex red assay of Y232X/R255L SpDAAO variants having positive and negative enrichment scores with D-Ala as a substrate (error bars indicate S.D of two independent measurements). Inset represents enzymatic activity of variants other than Y232. C) Amplex red assay of enriched Y232X/R255L SpDAAO variants against a range of amino acids (error bars indicate S.D of two independent measurements).

As expected, most of the R255L SpDAAO variants with positive enrichment scores showed considerable activity against D-Ala, while the variants with negative enrichment scores showed no reactivity against D-Ala, indicating that the negative enrichments identified using the SMART platform are true negatives. The same variants were also tested against other amino acids as substrates in the Amplex red assay and showed slight changes in the activity with different amino acids compared to the WT SpDAAO. These results highlight the importance of catalytic residue 232 for substrate specificity of SpDAAO (Fig. 3B).

In contrast to WT and R255L SpDAAO which accept a broad range of D-amino acids, the enriched variants (i.e. Y232S, Y232F, Y232I, Y232W) show reduced activity but distinct substrate specificity (Fig. 3C). The Y232S variant displayed decreased activity with D-Ala, but slightly increased activity with positively charged D-His and D-Arg, and hydrophobic D-Met, D-Leu and D-Phe. The Y223S variant of *Rhodotorula gracilis* DAAO (RgDAAO), a homolog of SpDAAO, has been reported to have 80-fold decreased catalytic turnover against D-Ala compared to the WT RgDAAO, a change caused by decreased substrate association and dissociation rates^[20b]^. This finding is in accordance with ours and supports the notion that the phenolic side chain of Tyr is essential for substrate binding. The enriched Y232F variant displays strong activity with D-Met and reduced activity with all other D-amino acids. This is in agreement with earlier findings on the pig kidney DAAO (pkDAAO) Y228F variant and RgDAAO Y223F variant which presented an overall decrease in catalytic turnover and increase in *K*_*M*_ indicating lower binding affinity for D-Ala^[20a]^. Interestingly, a subfamily-wide DAAO alignment carried out by the 3DM server (http://3dmcsis.systemsbiology.nl) indicates that besides Tyr at position 232 of SpDAAO which has a 90% conservation rate, F is present in part of the subfamily with 6.9% conservation rate (Fig. S13). The enriched Y232I variant exhibited an overall decline in activity, with some preference towards hydrophobic D-amino acids. Lastly, the Y232W variant demonstrated the highest activity against D-Leu and D-Met, and marginal activity against D-His, D-Arg and D-Phe. Notably, none of the tested SpDAAO variants shows considerable reactivity with L-Ala or the other L-amino acids, aligning well with previous observations^[20b]^.

The presence of Y232R, Y232P, and Y232D R255L SpDAAO variants (Fig. S14) in the enriched library is more complex to explain, given the loss of activity of these variants shown in the Amplex red assay (Fig. 3B, S15). Among other possibilities, we investigated the hypothesis of additional mutations in the regions of the SpDAAO gene other than position 232, which could restore the activity loss caused by the Y232 mutation. To test the hypothesis, we carried out Nanopore long-read sequencing of the Y232X/R255L SpDAAO input and enriched libraries. The data revealed the presence of additional mutations associated with Y232R, Y232P, and Y232D variants in the enriched library that are not present in the input library. All three variants (Y232R, Y232P, and Y232D) show a restored R255 sequence involved in the dimer formation. To verify the contribution of R255 to the enzymatic activity, we prepared genes of mutants that combine WT SpDAAO sequence R255, with either Y232R, Y232P, or Y232D mutations and tested their activity in Amplex red assay. The observed partial activity rescue (Fig. S16) indicates that R255 confers the activity disrupted by Y232 mutagenesis by an unknown mechanism, which could involve changes in the active site shape due to its sequence proximity to M248, a part of the active site core amino acid residues. More importantly, all enriched variants exhibited enzymatic activity in their true sequence, confirming that none were false positives. To the best of the authors’ knowledge, this study is the first to comprehensively analyze the impact of position 232 in SpDAAO on the enzyme’s activity and substrate specificity. Given that a ten-minute reaction time was set for the DAAO activity-based selection in the current study, implementing multiple selection rounds under more stringent conditions is anticipated to improve the enrichment of best-fit variants.

In summary, we have successfully developed the SMART platform for DAAO engineering, with apparent potential for the engineering of other oxidases of interest. Furthermore, SMART represents a universal *in vitro* enzyme engineering platform that can be easily customized for a range of enzymes with diverse chemistries by modifying the auxiliary unit components and enzyme activity-based selection conditions. Given the advantages of using commonly available reagents, a simple protocol, easy operation, short time, and efficient inclusion of functional variants in the enriched library, this platform holds the potential of becoming a valuable directed evolutionary tool for tailor-made enzyme development.

## METHODS

### 1.1 Preparation of pRSET-DAAO-R255L plasmid

pRSET-T26(QA) plasmid^[21]^ was used as the template for the preparation of the vector backbone by PCR with In-fusion_DAO(vec)_Fw and In-fusion_DAO(vec)_Rv primers. The plasmid pET21a-DAAO (obtained from Dr. T. Ito) was used as the template for the preparation of insert portion in PCR using InversePCR_DAOmono_R255L_Fw and InversePCR_DAOmono_R255L_Rv primers. PCR products after *Dpn*I digestion were purified using FastGene PCR/gel extraction kit (Nippon genetics) and assembled using HiFi DNA Assembly (New England Biolabs) at 5μl scale by 1-hour incubation of the mixture at 50°C. The resulting assembly was transformed into DH5α *E*.*coli* competent cells followed by plating on LB medium (with 100 μg/ml ampicillin), and incubation at 37°C overnight. Colonies with correct sequence were re-inoculated into 4mL liquid LB medium including ampicillin, and incubated overnight at 37°C. Plasmid extraction was conducted using Fastgene plasmid mini kit (Nippon Genetics): standard protocol.

### 1.2 Preparation of pRSET-DAAO-E49I, -W253I, -R255L, and -Y232A/R255L plasmids

The plasmids were prepared using the same method described in 1.1, with corresponding InversePCR_DAOmono primers for preparation of the insert portion.

### 1.3 Analysis of WT and R255L SpDAAO oligomeric state

The plasmids pET21a-DAAO and pET21a-DAAO-R255L were transformed into *E*.*coli* BL21(DE3) competent cells and cultured in liquid LB medium with 100 μg/ml ampicillin at 37°C until the OD reached 0.4-0.6. Then, IPTG was added at 0.1 mM to induce protein expression, and culture was continued at 16°C for 24 hours when the cells were harvested by centrifugation (15,000 rpm, 10 min, 4°C) and suspended in *E. coli*/Yeast Protein Extraction Buffer (TCI) according to the manufacturer’s protocol. The mixture was rotated at room temperature for 10 min. The suspension was then centrifuged (15,000 rpm, 10 min, 4°C), and the supernatant (soluble fraction) was used for the purification of SpDAAO variants by Ni-NTA affinity chromatography. Residual imidazole was removed by dialysis or filtration using a 10 kDa MWCO membrane (PALL corporation). After purification and imidazole removal, the enzyme samples were separated on the NativePAGE™ 4 to 16% gel (Thermo Fisher) at 150 V at room temperature for 90-120 min, using NativePAGE™ Running Buffer kit (Thermo Fisher). After electrophoresis, the gel was used for Western blotting and detection using Anti-His-tag mAb-HRP-DirecT (MBL Life Science) with ECL Prime (Cytiva) as the HRP substrate.

### 1.4 Cell-free protein synthesis (CFPS) of SpDAAO variants

Prepared pRSET plasmids with SpDAAO genes were used as the templates for PCR amplification with F1 and R1 primers followed by *Dpn*I digestion. About 50 ng of each purified template DNA was used in CFPS with Pure*frex* 2.1 kit (GeneFrontier) at 20 μl scale (Solution I, Solution II, Solution III, Cystein, and GSH: composition according to manufacturer’s protocol) with 20 mM FAD, a co-factor required for SpDAAO, and 0.8 μl of FluoroTech™ GreenLys (Promega) for the detection purposes. Samples were incubated at 37°C for 2 hours and centrifuged to separate soluble and insoluble fractions. Soluble fraction was used for SDS-PAGE analysis with fluorescence detection using an LED light imager equipped with a green band-pass filter and imaging software MISVS II (Biotools), and for enzyme activity assays.

### 1.5 4-aminoantipyrene assay of SpDAAO variants

4-Aminoantipyrene assay was used to evaluate the activity Y232X/R255L and WT Y232X input and enriched libraries. H_2_O_2_ released by DAAO reaction is used in consecutive HRP reaction with 4-Aminoantipyrene in the presence of phenol. The resulting quinoneimine shows absorption maximum at 505 nm with an extinction coefficient of 6.58mM^-1^cm^-1^. The reactions were set in a 96-well transparent plate (Thermo Fisher Scientific), with each well containing 5 μl of CFPS-made SpDAAO, 10 mM D-Ala, 50 μg of 4-aminoantipyrene, 25 μg of phenol and 2 U/mL HRP in 150 μl of 10 mM Tris HCl pH 8.0. The reaction mixture was incubated for 30 minutes at 37ºC and the absorbance was measured using the plate reader Infinite 200 PRO (Tecan). The standard line was prepared by using different concentrations of H_2_O_2_ (0, 0.1, 0.5, 5.0, 10 μM) in a reaction with 2 U/mL HRP, 25 μg of phenol, and 50 μg of 4-aminoantipyrene.

The activity was calculated using the following formula ^[22]^:

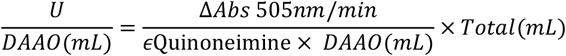

### 1.6 Amplex red assay of SpDAAO variants

Amplex red assay was used to evaluate the activity of all SpDAAO variants. H_2_O_2_ released by the DAAO reaction is used in consecutive HRP reaction with Amplex red as a substrate. The resulting resorufin shows fluorescence at 540/590 excitation/emission wavelength and absorbance at 570 nm. The reactions were set in a 96-well black plate (Thermo Fisher Scientific), with each well containing 1 μl of CFPS-made or displayed SpDAAO in 40 μl 0.25 M phosphate buffer pH 7.5, with 2.5 or 10 mM D-Ala as the substrate and 100 μl of a working solution composed of 0.1 mM Amplex red reagent, and 2 U/mL HRP in the same buffer. The fluorescence intensity was measured using the plate reader Infinite 200 PRO (Tecan) for 30-60 min continuously.

Due to better stability of detection, we later adopted absorbance detection. For this, we used the same reaction system in a Nunclon 96 Flat Bottom Transparent (Thermo Fisher) 96-well plate with continuous absorbance detection at 570 nm for 30 min. The standard line was prepared by using different concentrations of H_2_O_2_ (0, 0.1, 0.5, 5.0, 10 μM) in a reaction with 2 U/mL HRP and 0.1 mM Amplex red.

The activity was calculated using the following formula ^[22]^:

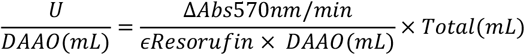

Where, *ϵResorufin* stands for the molar extinction coefficient of resorufin at 570 nm (54 mM^−1^ cm^−1^), *Total(mL)*is the final volume of the reaction mixture, *DAAO(mL)*is the volume of enzyme and *min* indicates the reaction time in minutes.

### 1.7 CFPS of APEX2-scCro fusion protein

pRSET-APEX2-scCro plasmid was used as the template in PCR with F1 and R1 primers followed by *Dpn*I digestion and PCR product purification. The resulting DNA was used as a template in CFPS using a PURE*frex*2.1 kit at 20 μl scale as described in 1.4, with 0.8 μl FluoroTech™ GreenLys and 10 μM hemin. The mixture was incubated at 37°C for 2 hours and centrifuged at 15000 rpm for 10 min at 4^º^C for the separation of soluble and insoluble fractions. Soluble fraction was analyzed by SDS-PAGE with fluorescence detection using an LED light imager.

### 1.8 Preparation of mRNA-displayed SpDAAO variants

pRSET-DAAO-R255L or pRSET-DAAO plasmid was used as a template in PCR with New Left and cnvK_New Ytag primers for preparation of DNA template for *in vitro* transcription by RiboMAX Large-scale RNA production system-T7 (Promega). The reaction mixture was incubated at 37°C for 2 h, followed by RNA purification using NucleoSpin RNA kit with on-column DNA digestion (Takara-Bio). The quality of synthesized RNA was estimated by Urea-PAGE after staining with SYBRGold, and the concentration was measured by absorbance at 260 nm using NanoDrop (Thermo Fisher). Gels were visualized by an LED light imager. Photocrosslinking of mRNA and cnvK puromycin linker^[14]^ (EME, Japan; Fig. S1) was done on a 20-μL scale by hybridization of the mRNA to the linker under the following thermal cycling protocol: 0.4°C/sec from 90°C to 70°C and 0.08 °C/sec from 70°C to 25°C, and subsequent UV irradiation at 365 nm for 4 min. Analysis of the crosslinked product was done by 8M Urea-PAGE with two detection methods, using fluorescent signal detection coming from the fluorescein attached to the linker, and staining with SYBRGold. The gels were visualized with an LED light imager. Crosslinked products were used for the *in vitro* translation to synthesize mRNA display using PURE*frex*2.1 kit on a 25-μL scale. Incubation of the mixture at 37°C for 30 min was followed by 5 min incubation at 37°C after addition of 20 mM EDTA to release the ribosomes. Analysis of mRNA display formation was done by SDS-PAGE after RNAse A treatment (incubation for 1 hour at 37°C) to digest the mRNA and reduce the complex size.

### 1.9 Preparation of single-molecule display

ORC hairpin linker is a single-stranded DNA of 81 bases including 19 complementary to the area of the mRNA outside of the DAAO gene. Folding of the hairpin structure was done on an 11.5 μl scale in a solution containing 1.97 mM of ORC DNA, 10 mM of Tris-HCl pH 8.00, 1 mM of EDTA and 1x Binding buffer (10 mM Tris-HCl buffer pH 8.0 with 1 mM EDTA, 1M NaCl, and 0.1% Tween 20) using the thermal cycling at 95°C:3min, 70°C:1min, 25°C:∞, with a ramp rate of 0.4°C/sec from step 1 to 2 and a ramp rate 0.1°C/sec from step 2 to 3.

Binding of the ORC hairpin linker to the SpDAAO mRNA display is performed by mixing the above ORC hairpin solution with 15 μl of SpDAAO mRNA display under the following conditions: 25°C:1 min, 40°C:1 min, 25°C:∞, with a ramp rate of 0.4°C/sec from step 1 to 2 and a ramp rate 0.1°C/sec from step 2 to 3.

The above mixture was then immobilized to the magnetic streptavidin microbeads (Dynabeads MyOne Streptavidin C1, Invitrogen) by mixing on a rotator for 30 min at 25°C. Immobilized complexes were washed with 1x Binding buffer and with 1x PBS. Beads were then mixed with 2.2 μl of CFPS-made APEX2-scCro, 2.2 μM hemin, and 41.8 μl of 1xPBS and incubated for 1 h at 4°C on a rotator. The beads with bound single-molecule display complexes were washed with 1xPBS.

Then the activity of tandem enzymatic reaction between SpDAAO and APEX2 was measured using the Amplex red assay as mentioned in passage 1.5.

The single-molecule display complexes were released from the beads by RNaseT1 treatment for 60 min at 30°C on a rotator. The collected supernatant (50 μl) containing single-molecule display complexes was separated into two fractions, 10 μl used for evaluation of the input library and the remaining 40 μl used for the activity-based selection.

### 1.10 Preparation of DNA templates for model library selection

Genes of R255L SpDAAO (active SpDAAO) and Y232A/R255L SpDAAO (inactive SpDAAO) were amplified by PCR using the corresponding plasmids as templates with New Left and cnvK_New Ytag primers. Amplified PCR products were column-purified and their concentration was evaluated by NanoDrop. Purified DNA was mixed in 1:1, 1:10, and 1:100 molar ratios of active and inactive SpDAAO genes respectively, to obtain the corresponding DNA model libraries. The libraries were used for preparation of the corresponding single-molecule display libraries according to 1.7 and 1.8.

### 1.11 DAAO activity-based selection

SpDAAO activity-based selection was performed on a 60 μl scale using 40 μl of single-molecule display solution. The reaction mixture was composed of 0.9 μl of 100X biotin-tyramide (Cosmo Bio), 1.6 μM hemin, 12.1 μl of 1X PBS, 7.5 mM D-Ala, and 0.6 mM FAD. The reaction mixture was incubated for 30 min at 37°C. In the next step, the unreacted biotin-tyramide was removed by ultrafiltration with a 10 kDa MWCO membrane (PALL corporation). The recovered upper residual liquid (~100 μL) was then immobilized onto the Streptavidin MyOne C1 magnetic microbeads from 20 μL suspension for 30 min at 25°C on a rotator. Then the mRNA display was converted to cDNA display with ReverTra Ace reverse transcriptase (Toyobo) at 42°C for 30 min in the presence of 1 μl of Recombinant RNAse inhibitor (Toyobo), 1mM of dNTP, and 100 U of ReverTra Ace enzyme for both enriched (biotinylated library after selection/ beads fraction) and input (original /library before selection) libraries.

### 1.12 Analysis of DNA recovered after model library selection

The input library, enriched library, and remaining pool (supernatant fraction) were used for PCR amplification with Nested_Fw1 and 2ndR DAAO_Rv1 primers in the first PCR, and with In-fusion_DAO_Fw and In-fusion_DAO_Rv primers in the second PCR with 1μL of the first PCR reaction mixture as a template. Five μL of the second PCR product was subjected to restriction digestion with *Xho*I and analyzed by agarose gel electrophoresis. The band pattern was compared with that of digested R255L SpDAAO and Y232A/R255L SpDAAO DNA to identify the enriched DNA.

### 1.13 Preparation and selection of Y232X libraries

The pRSET-DAAO-R255L plasmid and pREST-DAAO plasmids were used as templates for the inverse PCR with Y232(NNK)_Fw and Y232(NNK)_Rv primers, followed by *Dpn*I digestion and PCR product purification. About 50 ng of PCR product was used in Hifi assembly (New England Biolabs) with 2x HiFi master mix at 5 μl scale followed by PCR with New Left and cnvK_New Ytag primers to prepare the initial DNA libraries for *in vitro* transcription. The following steps (*in vitro* translation, single-molecule display preparation, activity-based selection, and DNA recovery) were done according to the protocols used for model libraries.

### 1.14 Analysis of the input and enriched Y232X SpDAAO libraries by next-generation sequencing

PCR-amplified cDNA pools of the enriched and input libraries were used for the preparation of 150 bp-long DNA fragments with codon of the 232 position in the middle of the fragment, by a PCR with NGS_DAAOcontrolLib_Fw1-6 and NGS_controlLib_Rv primers. Amplified fragments were gel-purified and analyzed by DNA-PAGE and Sanger sequencing to estimate the quality of preparation. Such prepared DNA was analyzed using the NextSeq550 Illumina platform in a single-read mode at the Center for Gene Research of Nagoya University.

To investigate the full gene length, libraries were analyzed by long-read Nanopore sequencing. PCR-amplified cDNA pools of the enriched and input libraries were used for another round of PCR amplification with NGS_InfusionDAAO 1-4 and Infusion DAAO_Rv primers. Amplified fragments were gel-purified and analyzed by DNA-PAGE and Sanger sequencing to estimate the quality of preparation. Prepared DNA was sequenced with a MinION Flongle Flow Cell (R10.4.1, Oxford Nanopore Technologies) on a MinION sequencing device (Mk1B) with a Flongle adapter following the manufacturer’s instruction. The basecalling was performed with MinKNOW software in real-time during the sequencing with Super Accurate (SUP) model.

### 1.15 Bioinformatics analysis of NGS data

The Illumina sequencing data were quality-filtered by SeqKit^[23]^. The quality threshold was set to Q20 and sequences that passed the criteria were used for further analysis. Each step was accompanied by testing the statistics of the analysis to ensure sufficient data were retained for the next step. In the following, only the DNA sequences corresponding to the SpDAAO gene were taken and converted into amino acid sequences using Bio-python. To exclude the unexpected outcomes, only the reads with sequence motif around position 232 (SDT*<u>X</u>*IIP, where X indicates sequence at 232) were retained for the analysis. The enrichment factor of each amino acid was calculated using the normalized count number of each residue in input and enriched libraries.

The Nanopore sequencing data was analyzed using the local Galaxy platform (https://galaxyproject.org). The fastq data were prepared for data processing using FASTQ Groomer (v1.1.5). After reverse-complementary processing of the reads with Manipulate FASTAQ (v1.1.5)^[24 a]]^, the reads were trimmed for remaining the DNA sequence from the start codon to the last amino acid of SpDAAO with Cutadapt^[24 b]]^(v4.0+galaxy1) using recognition sequences ATGACTAA and ATTGGCT, respectively. Finally, the mutation percentage calculation was performed with CRISPResso2^[24 c]]^ (v0.1.1).

### 1.16 Preparation of Y232X SpDAAO variants

The inverse PCR was done using pRSET-DAAO-R255L plasmid as the template with Y232(NNK)_Rv and each DAAO_Y232R_Fw, DAAO_Y232I_Fw, DAAO_Y232P_Fw, DAAO_Y232S_Fw, DAAO_Y232W_Fw, DAAO_Y232D_Fw primers with overlapping forward and reverse strands. The long linearized products were self-assembled using HiFi assembly. About 5 μl of the assembly mixture was used for the transformation of competent DH5α cells, which were plated on LB/ampicillin plates and incubated at 37°C for 17 hours. The colonies with correct sequences were inoculated in an LB medium and incubated overnight at 37°C for plasmid extraction with Fastgene plasmid mini kit. The same procedure was used for the preparation of dimeric Y232R, Y232P, and Y232D variants using pRSET-DAAO plasmid as the template. Obtained plasmids were PCR-amplified with F1 and R2 primers, column-purified and used as templates for CFPS using the PURE*frex*2.1 kit under the conditions described in 1.4.

## ACKNOWLEDGMENT

The authors are thankful to Dr. Takashi Kanamori from Gene Frontier, Japan for providing PURE*frex* components for cell-free protein synthesis, and Dr. Issey Takahashi from Nagoya University, Institute of Transformative Bio-Molecules for preparation of graphic materials.

