## Supporting information for "SMART platform for enzyme engineering: A novel single-molecule display approach for evolution and functional studies of D-amino acid oxidase"

[c] Dr. M. Camagna
Department of Plant Production Sciences
Nagoya University
Furo-cho, Chikusa Ward, Nagoya, Aichi 464-8601

[d] Professor N. Nemoto
Department of Applied Chemistry
Saitama University
255 Shimookubo-cho, Sakura Ward, Saitama, 338-8570

#### 1.5 4-aminoantipyrene assay of SpDAAO variants

4-Aminoantipyrene assay was used to evaluate the activity Y232X/R255L and WT Y232X input and enriched libraries. $H_2O_2$ released by DAAO reaction is used in consecutive HRP reaction with 4-Aminoantipyrene in the presence of phenol. The resulting quinoneimine shows absorption maximum at 505 nm with an extinction coefficient of 6.58$mM^{-1}cm^{-1}$. The reactions were set in a 96-well transparent plate (Thermo Fisher Scientific), with each well containing 5 µl of CFPS-made SpDAAO, 10 mM D-Ala, 136 µg of 4-aminoantipyrene, 63 µl of phenol and 2 U/mL HRP in 150 µl of 10 mM Tris HCl pH 8.0. The reaction mixture was incubated for 30 minutes at 37$^{o}$C, and the absorbance was measured using the plate reader Infinite 200 PRO (Tecan). The standard line was prepared by using different concentrations of commercially available pkDAAO (Sigma-Aldrich) in 0, 1.67, 3.33, 5.0, and 6.67 µg/mL concentrations, with other reaction components kept as described above.
The activity was calculated using the following formula [22]:

$$\frac{U}{DAAO(mL)} = \frac{2\ \times \Delta Abs\ 505nm/min}{\epsilon \text{Quinoneimine} \times\ DAAO(mL)} \times Total(mL)$$

#### 1.6 Amplex red assay of SpDAAO variants

Amplex red assay was used to evaluate the activity of all SpDAAO variants. $H_2O_2$ released by the DAAO reaction is used in consecutive HRP reaction with Amplex red as a substrate. The resulting resorufin shows fluorescence at 540/590 excitation/emission wavelength and absorbance at 570 nm. The reactions were set in a 96-well black plate (Thermo Fisher Scientific), with each well containing 1 µl of CFPS-made or displayed SpDAAO in 40 µl 0.25 M phosphate buffer pH 7.5, with 2.5 or 10 mM D-Ala as the substrate and 100 µl of a working solution composed of 0.1 mM Amplex red reagent, and 2 U/mL HRP in the same buffer. The fluorescence intensity was measured using the plate reader Infinite 200 PRO (Tecan) for 30-60 min continuously.
Due to better stability of detection, we later adopted absorbance detection. For this, we used the same reaction system in a Nunclon 96 Flat Bottom Transparent (Thermo Fisher) 96-well plate with continuous absorbance detection at 570 nm for 30 min. The standard line was prepared by using different concentrations of $H_2O_2$ (0, 10, 20, 30, 40 µM) in a reaction with 2 U/mL HRP and 0.1 mM Amplex red.
The activity was calculated using the following formula [22]:

$$\frac{U}{DAAO(mL)} = \frac{\Delta Abs570nm/min}{\epsilon Resorufin \times\ DAAO(mL)} \times Total(mL)$$

Where, $\epsilon Resorufin$ stands for the molar extinction coefficient of resorufin at 570 nm (54 $mM^{-1}$ $cm^{-1}$), $Total(mL)$ is the final volume of the reaction mixture, $DAAO(mL)$ is the volume of enzyme and $min$ indicates the reaction time in minutes.

#### Supporting Tables

Table S1. Count numbers and enrichment factors of SpDAAO variants in WT Y232X SpDAAO libraries.

| Residue at 232 position | Input library counts | Enriched library counts | Enrichment factor |
|---|---|---|---|
| A | 688554 | 530709 | 0.8856323 |
| F | 405187 | 354675 | 1.00579639 |
| V | 229641 | 233222 | 1.16695762 |
| L | 245242 | 229207 | 1.07391035 |
| S | 156280 | 223470 | 1.64305017 |
| K | 243375 | 215517 | 1.01751441 |
| W | 159903 | 133476 | 0.95913904 |
| Y | 35091 | 5212 | 0.17066468 |
| G | 11932 | 2235 | 0.21522825 |
| C | 11352 | 1856 | 0.18786271 |
| E | 1535 | 1308 | 0.97911649 |
| R | 782 | 1034 | 1.51931835 |
| T | 8048 | 838 | 0.11964403 |
| D | 696 | 720 | 1.18866166 |
| I | 11715 | 656 | 0.06434229 |
| M | 8355 | 505 | 0.06945123 |
| P | 6021 | 409 | 0.07805301 |
| N | 258 | 311 | 1.38508263 |
| H | 105 | 157 | 1.71808779 |
| Q | 24 | 96 | 4.59615843 |
| Total | 2224096 | 1935613 | |

Table S2. List of oligonucleotide primers used in the study.

| Name | **Sequence (5'-3')** | **Use** |
|---|---|---|
| New Left | GATCCCGCGAAATTAATACGACTCACTATAGGG | Preparation of DNA template for cDNA display |
| cnvK_New Ytag | TTTCCACGCCGCCCCCCGTCCT | |
| Nested_Fw1 | GGGAGACCACAACGGTTTCC | Amplification of DNA after selection |
| Nested_Rv1 | TTTCCCCGCCGCCCCC | |
| 2ndR_DAAO_Rv1 | ATGATGATGATGGCTGC | |
| In-fusion_DAO_Fw | TATGACTAAGGAAAATAAGCCAAGAGATAT | |
| In-fusion_DAO_Rv | AGCCAATTTGATTTTAGGAAGAGC | |
| F1 | atctcgatcccgcgaaattaatacg | Preparation of templates for CFPS |
| R1 | tccggatatagttcctcctttcag | |
| In-fusion_DAO(vec)_Rv | TTTTCCTTAGTCATATGTATATCTCCTTCTTAAAGTTAAACAAAA | Preparation of vector backbone for assembly |
| In-fusion_DAO(vec)_Fw | AAAATCAAATTGGCTGGTGGAGGTTCGGCCGGG | |
| InversePCR_DAOmono_R255L_ Fw | ATCGATCGTGAAATTCACCCTGAAG | Preparation of insert portions of monomeric DAAO plasmid constructs |
| InversePCR_DAOmono_R255L_ RV | GTTTCCTGGTTGCATGAAACC | |
| InversePCR_DAOmono_E49I_Fw | ATCTACACTTCCCCTTGGGCTGG | |
| InversePCR_DAOmono_E49I_Rv | TACAGAACGATCTTCAGGCGTATA | |
| InversePCR_DAOmono_W253I_Fw | ATCGATCGTGAAATTCACCCTGAAG | |
| InversePCR_DAOmono_W253I_Rv | GTTTCCTGGTTGCATGAAACC | |
| Y232(NNK)_Fw | GGCAAGAACTCTGATACCNNKATTATTCCTCGTCCC | Preparation of Y232NNK mutation |
| Y232(NNK)_Rv | TATGACTAAGGAAAATAAGCCAAGAGATAT | |
| ORC hairpin linker | GATCCtatcaccgcgggtgataGTACGTTTTTTCGTACtatcacccgcggtgataGGATCAAATGATGATGATGGCTGCCT | ORC hairpin linker |
| NGS_DAAOcontrolLib_Fw1 | CTGATATAATCATGTTGTTTTGGTTAAGGCTCCTC | Preparation of libraries for NGS |
| NGS_DAAOcontrolLib_Fw2 | CTACTAATCGCATGTTGTTTTGGTTAAGGCTCCTC | |
| NGS_DAAOcontrolLib_Fw3 | CATACTTTTTCATGTTGTTTTGGTTAAGGCTCCTC | |

| | | |
|---|---|---|
| NGS_DAAOcontrolLib_Fw4 | CAATTCATACCATGTTGTTTTGGTTAAGGCTCCTC | |
| NGS_DAAOcontrolLib_Rv | CGATCCCAGTTTCCTGGTTGCA | |
| DAAO_Y232R_Fw | GGCAAGAACTCTGATACCCGCATTATTCCTCGTCCC | Preparation of Y232 mutation to R, I, P, S, W, and D residues. |
| DAAO_Y232I_Fw | GGCAAGAACTCTGATACCATTATTATTCCTCGTCCC | |
| DAAO_Y232P_Fw | GGCAAGAACTCTGATACCCCGATTATTCCTCGTCCC | |
| DAAO_Y232S_Fw | GGCAAGAACTCTGATACCAGCATTATTCCTCGTCCC | |
| DAAO_Y232W_Fw | GGCAAGAACTCTGATACCTGGATTATTCCTCGTCCC | |
| DAAO_Y232D_Fw | GGCAAGAACTCTGATACCGATATTATTCCTCGTCCC | |

### Supporting Figures

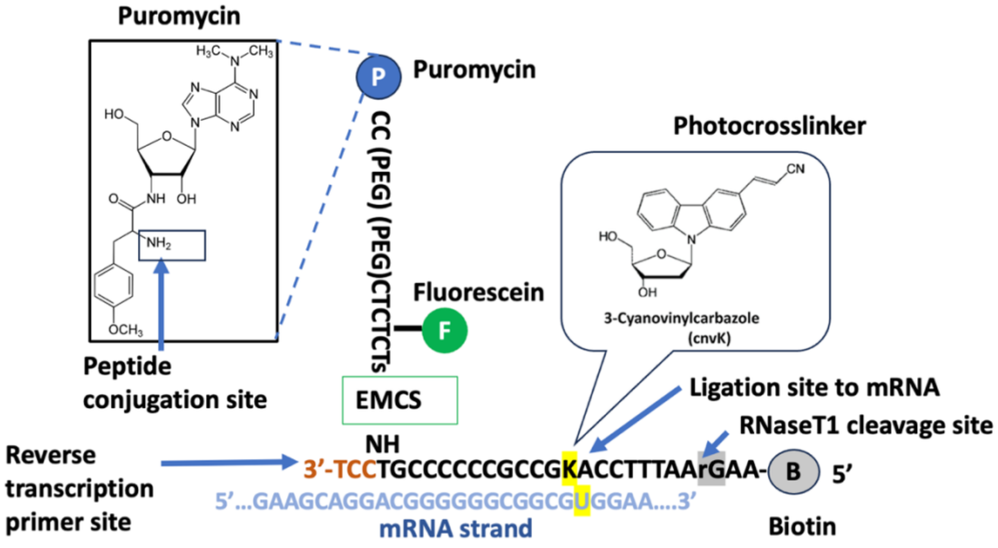

Figure S1. The schematic diagram of the puromycin linker. The site for mRNA photocrosslinking (cnvK), a biotin moiety for complex purification using streptavidin microbeads, and RNAseT1 restriction site for release of the complex from streptavidin microbeads are included in the scheme. Fluorescein is included for detection.

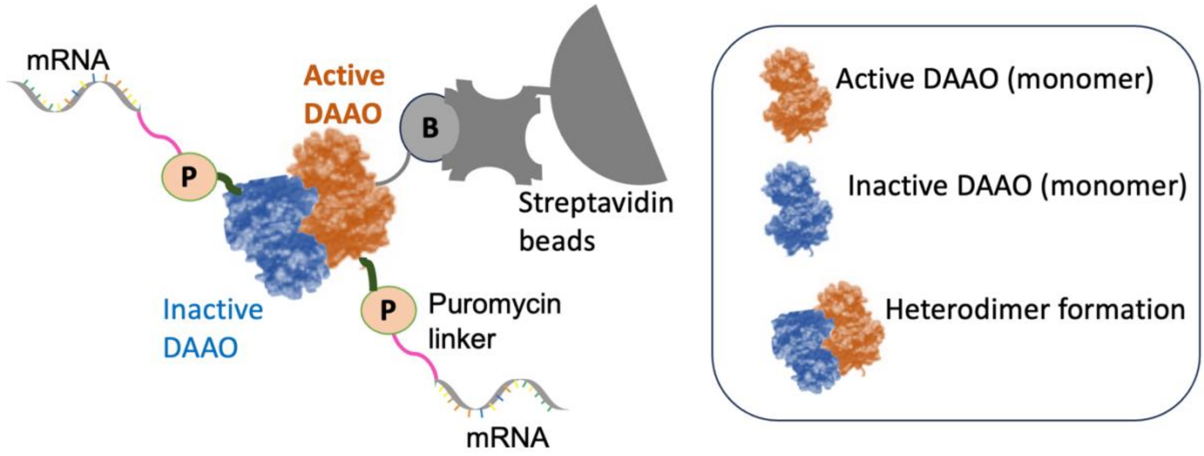

Figure S2. Schematic representation of theoretical SpDAAO heterodimer formation. Active SpDAAO variants are captured using the streptavidin beads pulldown. Heterodimer formation including an inactive variant could lead to false positive enrichment affecting the accuracy of the selection process.

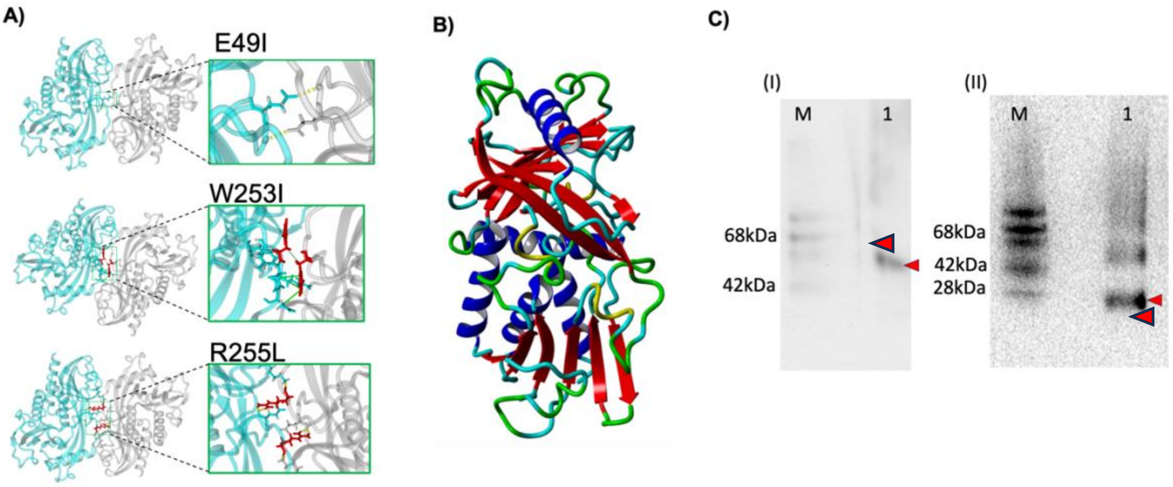

Figure S3. A) Model structure of SpDAAO dimer obtained from the Expasy Server (https://swissmodel.expasy.org). The amino acids identified as important for dimer formation are shown in stick representation (E49, W253, and R255). B) Model structure of R255L SpDAAO obtained by YASARA *Structure* and colored according to the secondary structure elements. C) Analysis of the R255L SpDAAO oligomeric state by Blue Native-PAGE. WT dimeric conformation is observed as a single band with mobility corresponding to 40-50 kDa, while R255L shows a pronounced band at around 28 kDa indicating the predominant monomeric structure.

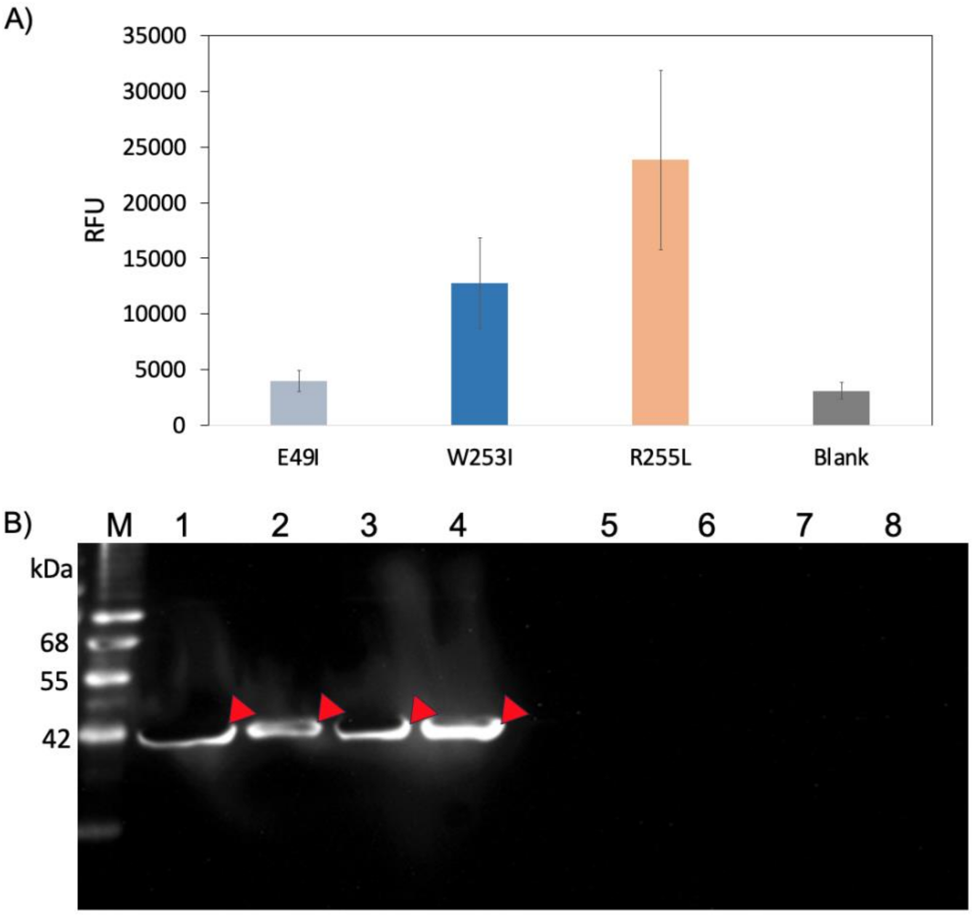

Figure S4. Evalution of E49I, W253I, and R255L SpDAAO variants. A) Amplex red assay of the SpDAAO variants with D-Ala as a substrate (error bars indicate S.D. of two independent measurements). B) Western blot analysis of CFPS-made SpDAAO variants detected using Anti-His-tag mAb-HRP-DirecT and ECL Prime HRP substrate. Lane M: Dr. Western marker, Lane 1: WT SpDAAO soluble fraction, Lane 2: E49I SpDAAO soluble fraction, Lane 3: W253I SpDAAO soluble fraction, Lane 4: R255L SpDAAO soluble fraction, Lane 5: WT SpDAAO insoluble fraction, Lane 6: E49I SpDAAO insoluble fraction, Lane 7: W253I SpDAAO insoluble fraction, Lane 8: R255L SpDAAO insoluble fraction.

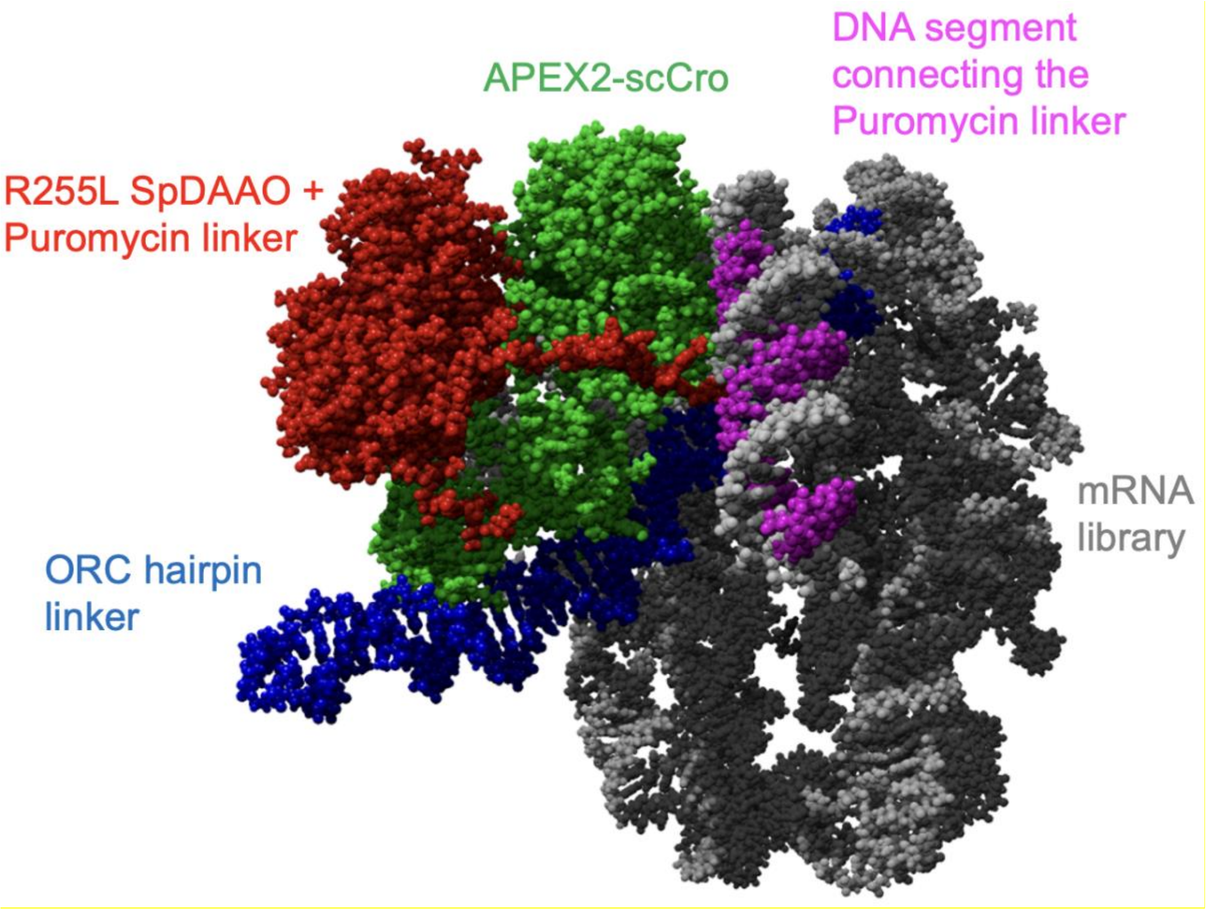

Figure S5. The model structure of R255L SpDAAO single-molecule display predicted by Alphafold3. mRNA (grey) is linked with the puromycin linker (connection showed in magenta) and R255L SpDAAO (red).

A)

ORC

| | | | | | | | | | | | | | | | | | | |
|---|---|---|---|---|---|---|---|---|---|---|---|---|---|---|---|---|---|---|
| T | A | T | C | A | C | C | | G | C | G | | G | G | T | G | A | T | A |
| A | T | A | G | T | G | G | | C | G | C | | C | C | A | C | T | A | T |

B)

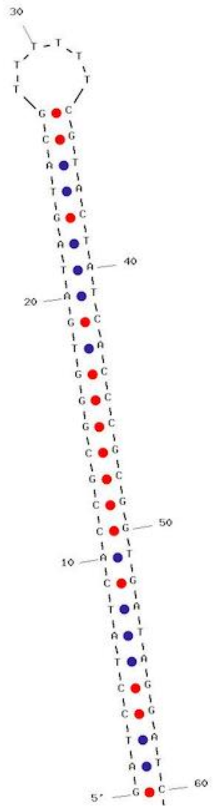

Figure S6. (A) Nucleotide sequence of the ORC hairpin DNA. B) The secondary structure of ORC hairpin DNA created using the IDT oligo analyzer tool (https://sg.idtdna.com/calc/analyzer).

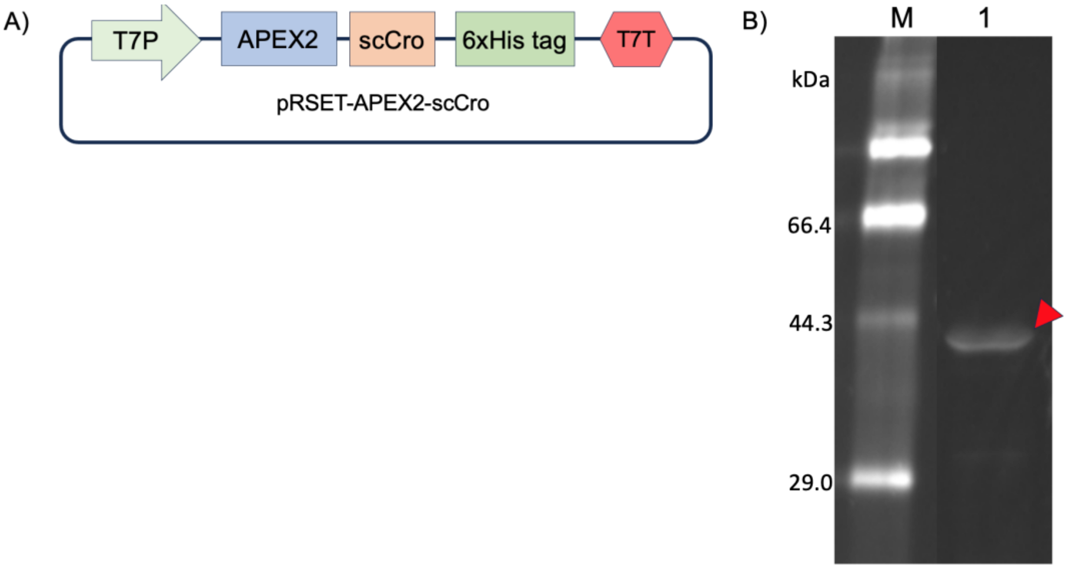

Figure S7. A) The scheme of the pRSET-APEX2-scCro plasmid. B) SDS-PAGE analysis of CFPS-made APEX2-scCro detected using FluoroTect GreenLys. Lane M: FITC marker, Lane 1: APEX2-scCro CFPS product (soluble fraction).

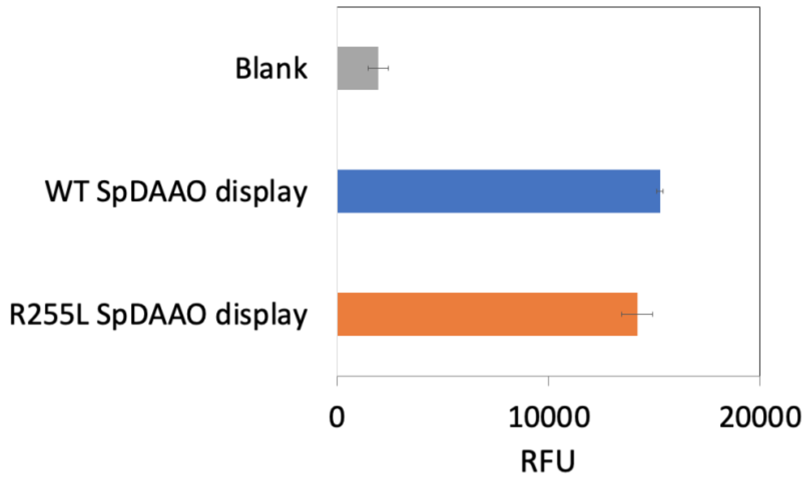

Figure S8. Amplex red assay of the tandem enzymatic reaction between displayed SpDAAO and APEX2 anchored to the mRNA display via scCro-ORC hairpin binding (error bars indicate S.D. of two independent measurements).

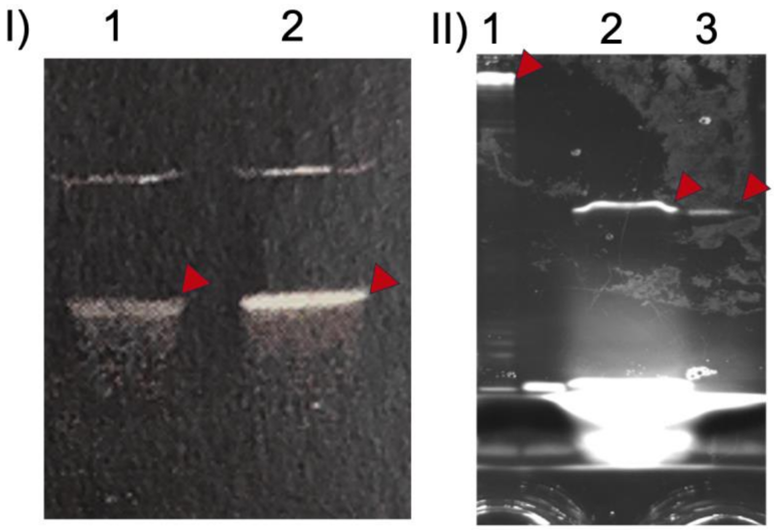

Figure S9. Evaluation of R255L SpDAAO display formation. (I) Urea-PAGE analysis of photocrosslinking reaction products. Lane 1: mRNA before photocrossliking, Lane 2: mRNA after photocrossliking; (II) SDS-PAGE analysis of mRNA display. Lane 1: mRNA after photocrossliking. Lane 2: mRNA display (soluble fraction) after RNAseA treatment for mRNA degradation, Lane 3: mRNA display (insoluble fraction) after RNAseA treatment for mRNA digestion. Arrows indicate the product bands.

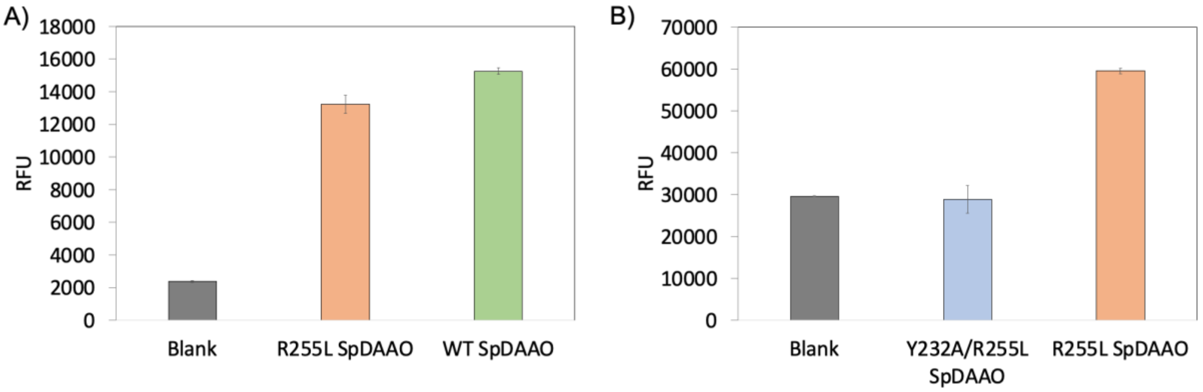

Figure S10. Amplex red assay of WT, R255L, and Y232A/R255L SpDAAO with D-Ala as a substrate. A) Evaluation of WT SpDAAO, R255L SpDAAO, and Y232A/R255L SpDAAO in mRNA displayed form. B) Evaluation of R255L monomeric SpDAAO, Y232A/R255L monomeric SpDAAO synthesized by CFPS (error bars indicate S.D. of two independent measurements).

| | | 230 | | |
|---|---|---|---|---|
| Consensus | Xaa | Xaa | Xaa | Xaa |
| Identity | | | | |
| | | 230 | | |
| 1. DAAO | Ser | Asp | Thr | Tyr |
| 2. Node362 | Glu | Thr | Thr | Tyr |
| 3. Node359 | Glu | Phe | Ala | His |
| 4. Node358 | Glu | Phe | Ala | His |
| 5. Node19 | Tyr | Ala | Thr | Tyr |
| 6. Node18 | Tyr | Glu | Thr | Tyr |
| 7. Node17 | Glu | Trp | Thr | Tyr |
| 8. Node16 | Glu | Phe | Thr | Tyr |
| 9. Node15 | Glu | Phe | Thr | Tyr |
| 10. Node14 | Glu | Phe | Thr | Tyr |
| 11. Node13 | Glu | Phe | Thr | Tyr |
| 12. Node12 | Thr | Phe | Thr | Tyr |
| 13. Node11 | Glu | Phe | Thr | Tyr |
| 14. Node10 | Thr | Trp | Thr | Tyr |
| 15. Node9 | Thr | Trp | Thr | Tyr |
| 16. Node8 | Thr | Trp | Thr | Tyr |
| 17. Node7 | Thr | Trp | Thr | Tyr |
| 18. Node6 | Thr | Trp | Thr | Tyr |
| 19. Node5 | Thr | Trp | Thr | Tyr |
| 20. Node4 | Thr | Trp | Thr | Tyr |
| 21. Node3 | Thr | Trp | Thr | Tyr |
| 22. Node2 | Asn | Ile | Ser | Tyr |
| 23. Node1 | Asn | Ile | Ser | Tyr |

Figure S11. Amino acid sequence alignment of predicted DAAO ancestral proteins with a focus on position 232 (as numbered in SpDAAO). Ancestral sequence reconstruction was performed using 1049 DAAO variants across 486 geni of fungal species.

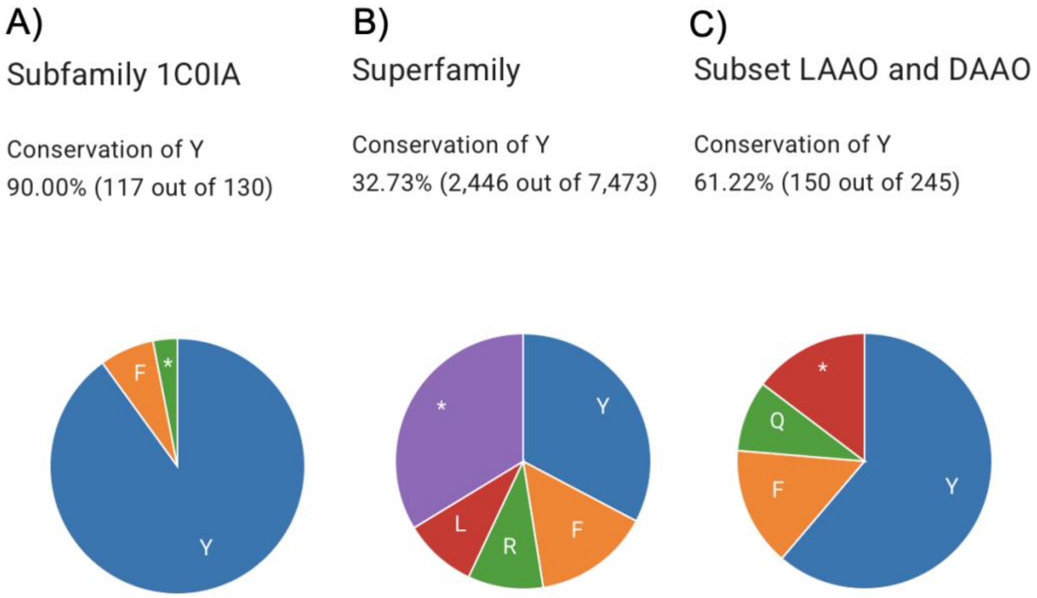

Figure S12. Pie charts showing the composition of amino acid sequences at 232 position of SpDAAO obtained from 3DM (http://3dmcsis.systemsbiology.nl) alignment. A) Alignment result according to the subfamily level revealing Tyr to be conserved at 90% sequences. The asterisk indicates other minor sequences (Cys, His, and Asn). B) Alignment result at the superfamily level revealing Tyr to be conserved at 32.7% sequences. The asterisk indicates other sequences. C) Alignment result at the subset of L-amino acid oxidases and DAAO indicating the presence of Tyr in about 61%, Phe in about 15%, and Gln in 8.9%. The asterisk indicates other sequences.

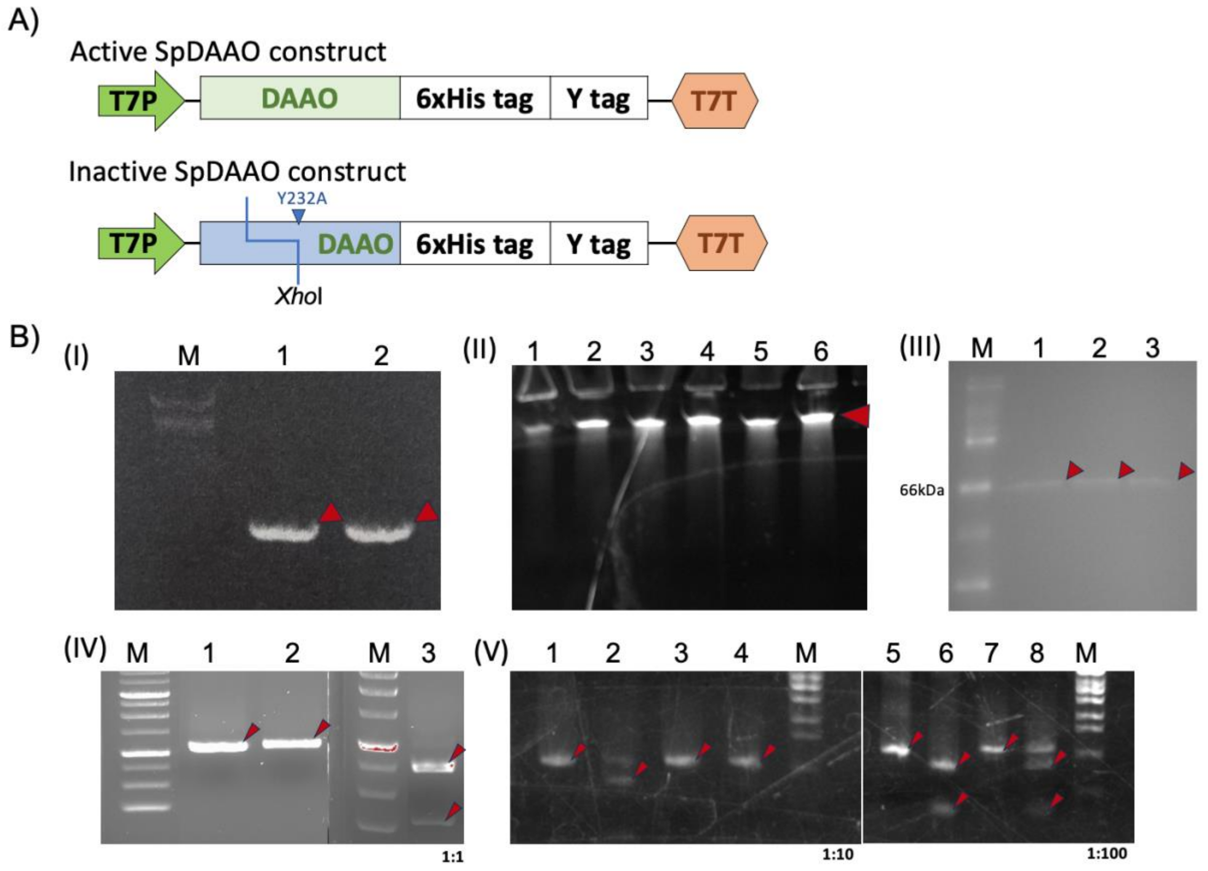

Figure S13. Evaluation of 1:1, 1:10, and 1:100 model library selection. A) The outline of the R255L SpDAAO active DNA construct and Y232A/R255L SpDAAO inactive DNA construct containing *Xho*I restriction site. B) (I) Agarose gel electrophoresis of DNA libraries. Lane M: λ-Eco T14 I digest, Lane 1: R255L SpDAAO DNA, Lane 2: Y232A/R255L SpDAAO DNA. (II) Urea-PAGE of photocrosslinked mRNA libraries after SYBRGold staining. Lane 1: 1:1 model mRNA library before photocrosslinking. Lane 2: 1:1 model mRNA library after photocrosslinking. Lane 3: 1:10 model mRNA library before photocrosslinking, Lane 4: 1:10 model mRNA library after photocrosslinking. Lane 5: 1:100 model mRNA library before photocrosslinking. Lane 6: 1:100 model mRNA library after photocrosslinking. (III) SDS-PAGE analysis of the mRNA display. Lane M: FITC marker (prepared in-house), Lane 1: 1:1 model library mRNA display, Lane 2: 1:10 model library mRNA display, Lane 3: 1:100 model library mRNA display. (IV) Agarose gel electrophoresis of PCR-amplified enriched and remaining libraries. Lane M: 1kbp DNA marker, Lane 1: 1:1 enriched library before *Xho*I treatment, Lane 2: 1:1 enriched library after *Xho*I treatment. Lane 3: 1:1 remaining library after *Xho*I treatment. (V) Agarose gel electrophoresis of PCR-amplified enriched and remaining libraries. Lane 1: 1:10 remaining library before *Xho*I treatment. Lane 2: 1:10 remaining library after *Xho*I treatment. Lane 3: 1:10 enriched library before *Xho*I treatment. Lane 4: 1:10 enriched library after *Xho*I treatment. Lane M: λ-Eco T14 I digest. Lane 5: 1:100 remaining library before *Xho*I treatment. Lane 6: 1:100 remaining library after *Xho*I treatment. Lane 7: 1:100 enriched library before *Xho*I treatment. Lane 8: 1:100 enriched library after *Xho*I treatment.

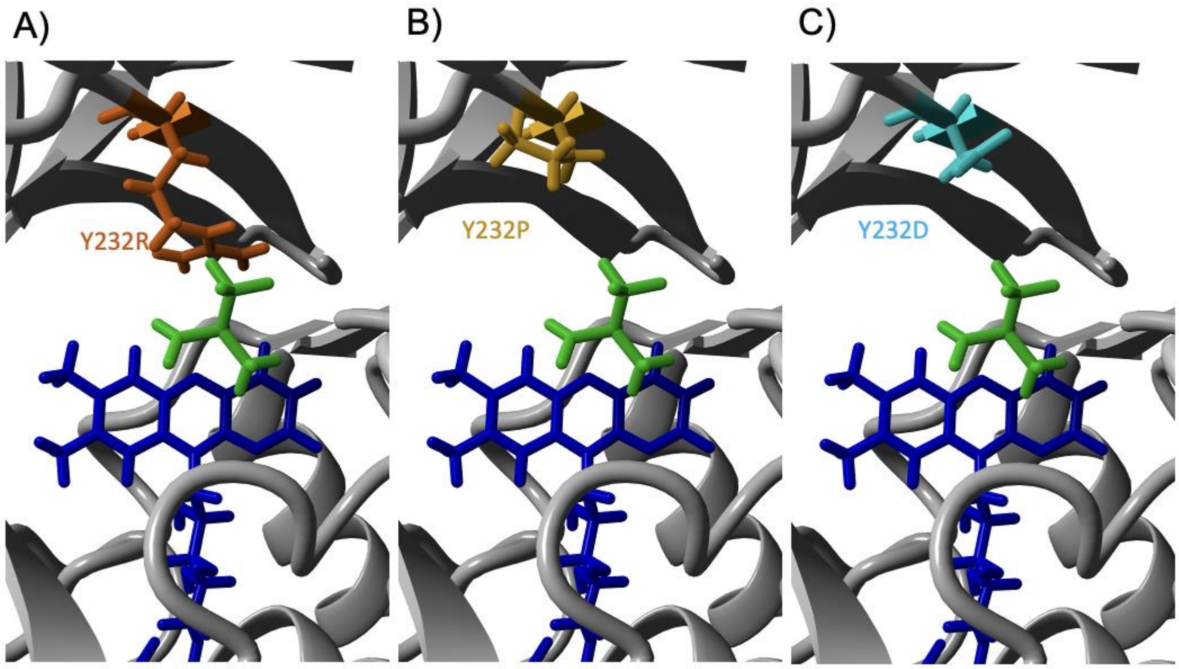

Figure S14. Model structures of Y232R/, Y232P/, and Y232D/R255L SpDAAO variants focusing on the active site. A) Y232R variant. B) Y232P variant. C) Y232D variant. FAD is colored blue and trifluoroalanine is colored green. Trifluoroalanine is adopted from the DAAO structure of *Rhodotorula toruloides* (PDB code: 1C0L).

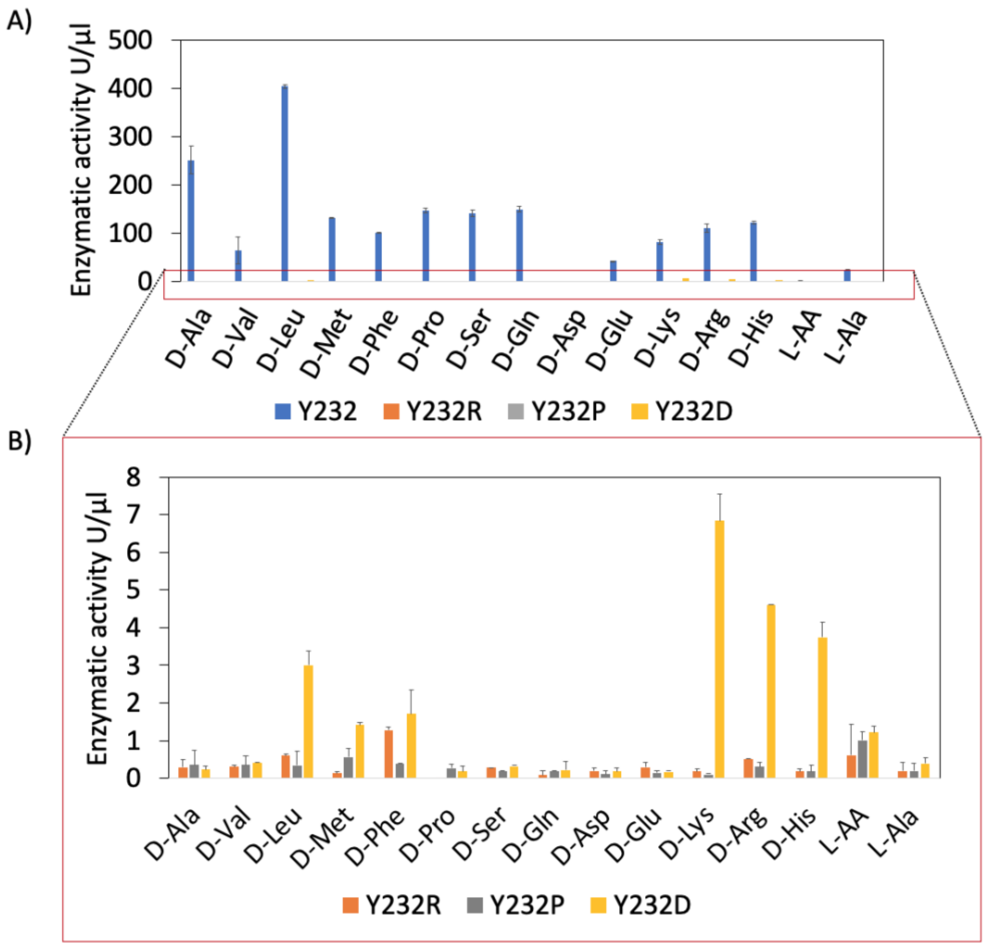

Figure S15. Amplex red assay of Y232R/, Y232P/, and Y232D/R255L SpDAAO variants against different amino acids as substrates. A) Activity of CFPS-made SpDAAO variants. B) Enlarged and rescaled activity of CFPS-made SpDAAO variants (error bars indicate S.D. of two independent measurements).

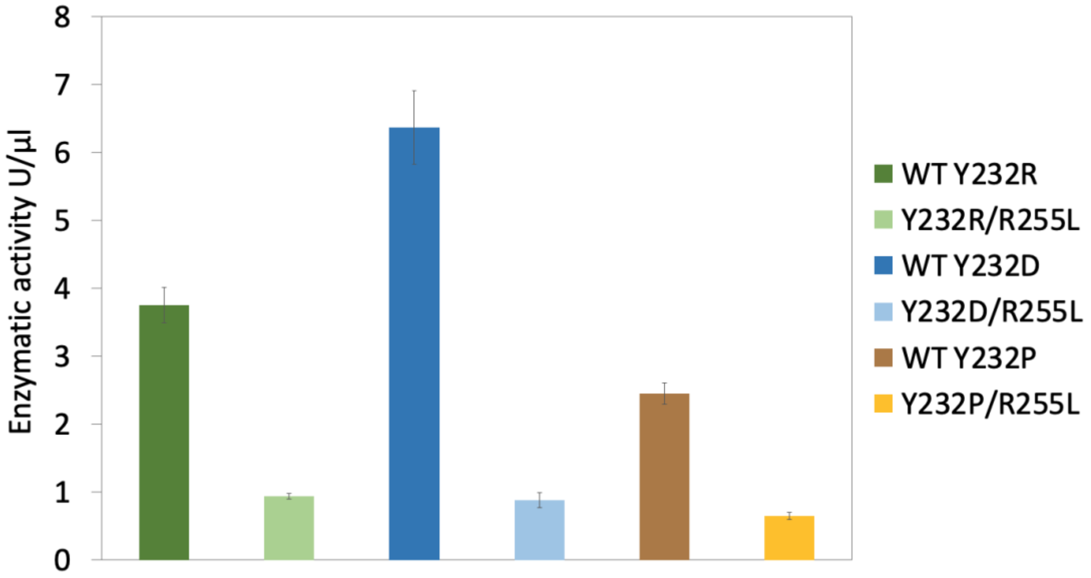

Figure S16. Amplex red assay of Y232R, Y232D and Y232P WT SpDAAO variants in comparison with R255L SpDAAO variants against D-Ala as the substrate (error bars indicate S.D. of three independent measurements).
